# Beyond one-size-fits-all: single-cell transcriptomic signatures predict drug efficacy and reveal responder subgroups in endometriosis

**DOI:** 10.64898/2025.12.09.693218

**Authors:** Raúl Pérez-Moraga, Cemsel Bafligil, Sarah Harden, Santiago García-Martín, María José Jiménez-Santos, Ioanna Tiniakou, Sophie Ribeiro-Volturo, Line Gies, María Teresa Pérez Zaballos, Cristina Fernández-Molina

**Affiliations:** endogene.bio, Paris, France; Computational Biomedicine Laboratory, Príncipe Felipe Research Center (CIPF), Valencia, Spain; Azura Biosciences, Madrid, Spain

## Abstract

Endometriosis affects ∼10% of reproductive-age women, yet targeted non-hormonal therapies remain unavailable, and treatment response is highly variable. Here, we apply a single-cell framework to resolve therapeutic heterogeneity at a resolution previously unattained in drug development efforts.

Using scRNA-seq profiles from eutopic and ectopic tissues, combined with a machine learning-based drug response model, we identified compounds predicted to revert disease-associated transcriptional states and map cell-type-specific vulnerabilities across patients and tissues. Our analysis revealed pronounced tissue-specific and inter-patient heterogeneity in predicted responses. Stromal, endothelial, and stem cell populations emerged as the dominant therapeutic targets, collectively revealing selective sensitivity to two recurrent drug classes, histone deacetylase and tubulin polymerisation inhibitors. Transcriptomic comparison of predicted responders and non-responders to these drugs pointed to conserved molecular programmes involving extracellular matrix remodelling, angiogenesis, and proliferative activation. These signatures were shared between eutopic and ectopic stromal compartments, supporting the feasibility of assessing therapeutic response using readily accessible eutopic tissue.

Our findings show that this single-cell framework can dissect therapeutic heterogeneity in endometriosis, support the development of precision non-hormonal therapies and identify responder subgroups relevant for patient stratification. Together, these results highlight that underlying molecular diversity in endometriosis necessitates therapeutic approaches beyond a one-size-fits-all model.

## Introduction

Endometriosis, an oestrogen-dependent chronic disease, impacts nearly one in ten women during their reproductive years and is characterised by the presence of endometrial-like cells outside the uterus, most commonly on the ovaries, pelvic peritoneum, and other abdominal organs (T K Saunders and Horne, 2025). It is a leading cause of debilitating pelvic pain, dysmenorrhea, and infertility (Elizur *et al*., 2025), substantially reducing the quality of life. Despite its high prevalence, endometriosis remains widely underdiagnosed, with an average diagnostic delay of 7 to 10 years (Fryer, Mason-Jones and Woodward, 2024), often preceded by multiple misdiagnoses (Bontempo and Schiff, 2025). Current treatments are predominantly palliative, relying on hormonal suppression or surgical excision, both associated with side effects, and recurrence, and limited long-term efficacy (Liao, Monsivais and Matzuk, 2024). Moreover, clinical responses to hormonal therapy are highly variable (Becker *et al*., 2017). For instance, Dienogest provides satisfactory outcomes in only about half to two thirds of patients, while those with advanced disease or concurrent adenomyosis frequently experience poor response (Nirgianakis *et al*., 2021; C.-H. Chen *et al*., 2025). This clinical variability in drug response is, at least in part, driven by inter-patient heterogeneity at the molecular and cellular level. Precision approaches capable of linking molecular profiles to therapeutic response are essential for addressing this heterogeneity and for developing effective, stratified and durable therapies for endometriosis.

An important translational application for this approach lies in its potential to inform clinical trial design. By stratifying patients according to predicted cell-type-specific sensitivities, these methods could enable more precise enrolment criteria, ensuring that participants most likely to benefit from a given therapy are represented. Additionally, they can serve as exploratory endpoints to interpret heterogeneous responses at trial completion. This may help address the high rate of clinical trial failure observed for endometriosis, where most completed trials have not published their results and are presumed to have failed (Guo and Groothuis, 2018). We hypothesize that heterogenous lesions and variable treatment responses confound outcomes. In oncology, both bulk and single-cell transcriptomics have been explored to predict response to treatment (Chen *et al*., 2022; Mahdi-Esferizi *et al*., 2024). A recent retrospective analysis found that cell-type specificity and disease-associated overexpression, derived from scRNA-seq data, are strong predictors of drug target success in clinical development (Dann *et al*., 2024). The authors noted that targets supported by scRNA-seq evidence could nearly triple the likelihood of a therapeutic advancing to Phase III clinical trials. Identifying predictive biomarkers of treatment response can, therefore, guide patient selection, enable rational drug repurposing and inform combination therapy design. Applying similar approaches to endometriosis may clarify why only some patients respond to a given therapy, reveal mechanisms of resistance and uncover biomarkers to monitor therapeutic efficacy.

The clinical and biological complexity of endometriosis presents major barriers to therapeutic advancement. Lesions vary widely in location, morphology, and molecular profile with multiple subtypes: ovarian endometrioma, superficial peritoneal lesions and deep endometriosis, often co-existing within a single individual (Alimi *et al*., 2018). Roughly 60-80% of lesions contain both epithelial and stromal cells, while the rest are stromal-dominant or heavily fibrotic (Nezhat *et al*., 2025). This anatomical and cellular heterogeneity is mirrored by molecular and functional differences in stromal, epithelial, endothelial and immune cell populations, each contributing distinct functional roles in angiogenesis, inflammation and lesion progression. Endometrial mesenchymal stem cells (eMSCs) and stromal cells (EnSC), in particular, are implicated in lesion establishment and persistence (Mariadas, Chen and Chen, 2025), while immune cells such as M2-polarised macrophages (Dai *et al*., 2025) and T cells (Knez, Kovačič and Goropevšek, 2024) contribute to chronic inflammation and tissue remodelling (Giudice, Liu and Irwin, 2025). These cellular and tissue-level variations likely underpin the heterogeneous drug responses observed in patients, highlighting the importance to characterise endometriosis at the molecular and cellular level.

Conventional bulk transcriptomics has advanced our understanding of endometriosis biology, but cannot resolve the contributions of individual cell types, including rare yet functionally important populations such as eMSCs, which constitute only ∼1% of endometrial cells (Gargett and Masuda, 2010). In bulk datasets, signals from distinct populations are combined (Saare *et al*., 2017), obscuring the specific roles that each cell type plays in disease progression and in shaping response to therapy. In addition, animal models of endometriosis fail to capture the full spectrum of human hormonal regulation, further limiting translational relevance (Malvezzi *et al*., 2020). Although transcriptomics-based drug repurposing efforts have yielded candidate compounds (Oskotsky *et al*., 2024, 2025), these studies relied on bulk data and non-primate models, restricting their ability to predict cell-type specific responses. Altogether, these limitations highlight the need for human-based, cell-type-resolved approaches to drug discovery.

scRNA-seq enables high-resolution dissection of transcriptionally distinct cell populations, dynamic states, and intercellular communication networks (Giudice, Liu and Irwin, 2025; Liu *et al*., 2025). This level of resolution is critical for identifying cell-type-specific drivers of disease and therapeutic vulnerability, particularly in conditions as heterogenous and complex as endometriosis (Ma *et al*., 2021). Computational tools such as *scTherapy* (Ianevski *et al*., 2024) integrate single-cell expression profiles with large-scale drug perturbation (Stathias *et al*., 2020) and dose-response datasets (Smirnov *et al*., 2018) to predict drug sensitivities at the single-cell level. Originally developed for oncology (Ianevski *et al*., 2024; X. Chen *et al*., 2025; Yates *et al*., 2025), these methods are applicable to endometriosis because both diseases are characterised by uncontrolled cell proliferation, aberrant survival signalling, and tissue invasion, with distinct subpopulations driving disease progression. Our recent work on menstrual blood-derived stem cells showed that early disease-associated epigenetic alterations converge on the same proliferative and pro-invasive pathways implicated in cancer biology (Tiniakou *et al*., 2025). Beyond the pathological similarities, compelling evidence connects endometriosis to the aetiology of both ovarian and endometrial cancers (Painter *et al*., 2018; Iyshwarya, Mohammed and Veerabathiran, 2021). Therefore, scTherapy is well-suited to explore therapeutic vulnerabilities in endometriosis.

Since eutopic endometrium represents both a source of ectopic seeding and a contributor to ongoing symptoms, assessing whether therapeutic response patterns are conserved between eutopic and ectopic tissues is essential. Such analyses could not only validate the robustness of predicted compounds but also establish eutopic stromal cells as a tractable model for preclinical evaluation. Demonstrating that therapeutic response biomarkers identified in ectopic lesions are conserved in the eutopic endometrium would furthermore enable their non-invasive interrogation via menstrual blood, providing a practical avenue for longitudinal monitoring and therapeutic assessment.

Here, we systematically evaluate the therapeutic landscape of endometriosis across ectopic lesions and eutopic endometrium and major non-immune populations. By integrating publicly available scRNA-seq datasets with large-scale compound perturbation profiles we predict drug sensitivities across key stromal, epithelial and endothelial populations, and reveal substantial heterogeneity in response both among patients and among cell types. Our analyses further link these predicted responses to underlying molecular programmes highlighting potential mechanisms of drug sensitivity and resistance. Finally, we show that predicted responder profiles are conserved between eutopic and ectopic sites, supporting the potential for non-invasive monitoring of therapeutic biomarkers. These findings provide a framework for cell-type-informed therapeutic prioritisation in endometriosis and establish a resource for advancing precision medicine in a disease long constrained by diagnostic and therapeutic stagnation.

## Materials and Methods

### scRNA-seq dataset acquisition and inclusion criteria

We systematically surveyed Gene Expression Omnibus (GEO) and Sequence Read Archive (SRA) for human endometriosis scRNA-seq datasets published between 2010 and 2023, using the search string: (endometriosis OR “endometrial disease”) AND ((”single cell” OR scRNA-seq OR “single-cell RNA sequencing”) AND human[Organism]). Inclusion criteria were: i) human samples from ectopic endometriosis, ovary, or eutopic endometrium; ii) availability of metadata (age, lesion type/stage, hormonal treatment); iv) raw sequence data in FASTA format); iv) 10X genomics technology to minimise platform batch effects. The final cohort comprised 110 samples from four datasets: GSE179640 (Tan *et al*., 2022), GSE213216 (Fonseca *et al*., 2023), GSE214411 (X. Huang *et al*., 2023), and PRJNA932195 (Shin *et al*., 2023), including eutopic and ectopic tissues from endometriosis patients and controls. Metadata were retained for stratified analyses (Supplementary Figure 1 and Supplementary Table 1).

### Data processing and quality control

Raw FASTA sequences were downloaded using sra-tools (3.0.10) toolkit and processed with CellRanger (v7.2) to demultiplex, align, and generate count matrices, including intronic reads. Up to 20,000 droplets per sample were included to recover low abundance cells and improve ambient RNA removal using CellBender (v0.3.0). Quality control (QC) excluded droplets with < 500 detected features or > 25% mitochondrial content, and potential doublets were removed using scDblFinder (v1.20). After filtering, 672,590 high-quality cells were retained for downstream analyses.

### Cell annotation using the Human Endometrial Cell Atlas

To harmonise cellular labels across datasets, we annotated cells at a principal lineage level using the Human Endometrial Cell Atlas (HECA) (Marečková *et al*., 2024). A support vector machine (SVM) classifier trained on 36 curated HECA cell types (25% held-out test set) achieved 91% accuracy (Supplementary Table 2). HECA labels were collapsed into major lineages (stromal, epithelial, endothelial, immune, smooth muscle, perivascular, fibroblast) for downstream analysis (Supplementary Table 3).

eMSC were not included among the predicted labels from the HECA-based SVM classifier. Therefore, we annotated this population *a posteriori* using canonical MSC surface markers Cells co-expressing CD44, CD73 (*NT5E*), and CD90 (*THY1*) with SCT-normalised expression ≥ 0.5 were classified as eMSCs. To ensure specificity and exclude epithelial stem-like populations, *SOX9* expression was used as a negative control. Only cells with low or absent *SOX9* expression were retained in the final eMSC annotation (n = 16,814).

### Pseudobulk matrix construction and DGE analysis

Pseudobulk gene expression matrices were generated by aggregating UMI counts per individual, lesion site, and main cell type, preserving inter-person variability. For the purposes of this analysis, anatomical sites were grouped into three tissue categories (Supplementary Table 4). Stromal cells were further grouped by menstrual cycle phase (proliferative n = 43,817, secretory n = 64,271 and exogenous hormone-treated n=15,139). Stromal cells from other menstrual phases were excluded from downstream analyses due to insufficient representation. Differential gene expression (DGE) analyses were performed using *edgeR* (v 4.0.16) on pseudobulk counts, controlling for dataset origin and tissue. Only samples with ≥ 25 cells and ≥ 50,000 reads were included. Both global (all tissues) and tissue-specific comparisons were conducted, with dataset included as a covariate. Counts were normalised using the trimmed mean of M-values method (TMM) and adjusted for multiple testing using the Benjamini-Hochberg method.

### Functional Enrichment

Exclusive DEGs for each cell type obtained at the global level were subjected to pre-ranked GSEA using the *fgsea* function (parameters: minSize = 15, maxSize = 500, nPermSimple = 10000) from the fgsea package, with the Hallmark v2024.1 collection as reference (Liberzon *et al*., 2015). We performed the functional enrichment analysis and selected the top 50 significantly upregulated genes (FDR < 0.05) per cell type based on logFC. The formula accounted for conditions, dataset, and tissue location:

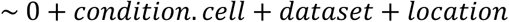

### Drug Prediction with scTherapy

We applied scTherapy to predict individual- and cell-type-specific responses across stromal cells, endothelial cells, and eMSCs. Immune populations were excluded due to limitations in current reference perturbation datasets. Pseudobulk DEGs for each individual, tissue, and cell type were used to query a large-scale drug perturbation database. High confidence compounds were prioritised based on predicted response score, absence of toxicity annotations, targeted mechanism, coverage across cell types or tissues, and emphasis on stem cell targeting. Known relevant compounds (ABT-751, proteasome inhibitors, HDAC inhibitors) were retained regardless of rank (Wang *et al*., 2022; Psilopatis *et al*., 2023; Wróbel *et al*., 2023).

### Aggregation of individualised predictions and clustering

For each cell population, a binary matrix indicated whether each of the top 10 predicted drugs was prioritised per individual, tissue, and cell type. Hierarchical clustering (*ward.D2*) was performed on binary dissimilarity matrices to identify shared drug response patterns. Heatmaps with dendrograms visualised patient stratification and response clusters, thereby identifying therapeutic clusters among patients.

### Transcriptomic drug response

To obtain the transcriptomic signatures of the different cell types of interest between inferred high responders and non-responders, stratified by cell type and lesion type, we subset the dataset to perform differential gene expression analysis using the *FindMarkers* function from the *Seurat* package. The *MAST* method was employed, including the dataset of origin as a covariate in the model (Finak *et al*., 2015). Genes with an FDR < 0.05 and an absolute logFC > 0.5 were considered significant.

### Over-representation analysis

Over-representation analysis (ORA) was conducted using only the genes upregulated in the high responder group for each of the drugs analysed. The ORA was performed with the *WebGestaltR* (v0.4.6) package (Elizarraras *et al*., 2024). Gene Ontology (GO) Biological Process terms were used as the ontological framework, with minimum and maximum category sizes set to 10 and 500, respectively. All genes annotated in the hg38 genome assembly served as the reference set, and upregulated genes in the high response groups were used as the input list. Statistical significance was determined using FDR-adjusted *p*-values, with a threshold of 0.05.

### Stemness score calculation

For the calculation of the stemness score for the cell types of interest, we used the R package *CytoTRACE2* (Kang *et al*., 2025). Specifically, we applied the function provided in the package with the parameters *batch_size* = 10000 and *smooth_batch_size* = 4000, since our datasets contained more than 10,000 cells. The function returns a vector of scores for each evaluated cell, ranging from 0 to 1, where lower values indicate more differentiated cells and higher values indicate more totipotent cells. We used a two-sided Kolmogorov–Smirnov tests and Cliff’s delta effect size estimates to determine if stemness scores differed significantly between responder groups.

### Correlation between cellular populations at transcriptomic level

With the aim of testing the similarity between stromal cells located in peritoneal lesions and those from the eutopic endometrium across different menstrual cycle phases, we generated a consensus transcriptomic signature. Specifically, we created a vector containing the 4,000 most variable genes, selected using the *FindVariableFeatures()* function from the Seurat package. We then generated pseudobulk profiles of stromal cells for each tissue and computed the Pearson distances between the resulting vectors.

Finally, to evaluate the concordance between transcriptional signatures from peritoneal lesions and ectopic endometrium, we compared their differential expression profiles using a hypergeometric test to assess the significance of overlapping DEGs and the Jaccard index to quantify similarity between up- and downregulated gene sets.

## Results

### High-quality single-cell atlas identifies both major and rare endometrial cell populations in endometriosis

To enable a unified analysis of endometriosis at the cell-type level across tissues and individuals, we first integrated and harmonised publicly available scRNA-seq datasets from eutopic and ectopic sites (GSE179640, GSE213216, GSE214411, PRJNA932195). This integrated dataset served as the basis for all downstream drug prediction analyses; it comprised 672,590 cells across 110 samples with metadata including age, menstrual cycle phase, hormonal treatment, and disease stage where available.

Cell annotation was performed in an agnostic manner by applying a support vector machine (SVM) classifier trained on the state-of-the-art endometrium atlas, the Human Endometrium Cellular Atlas (HECA) (Marečková *et al*., 2024). To assess the model’s reliability, a portion of the data was excluded from training and solely used for testing. The model correctly classified 91% of these cells. For interpretability, HECA labels were collapsed into seven broad cell lineages: stromal, epithelial, endothelial, immune, smooth muscle, perivascular, and fibroblast. A total of 506,165 high quality labelled cells were used for subsequent analysis (Figure 1A). Notably, eMSCs were not reliably captured by the classifier. This likely reflects that eMSCs are relatively rare and less well-characterised transcriptionally compared to major endometrial cell types, resulting in less distinct signatures in the reference atlas. Instead, a targeted, marker-based annotation strategy was applied using canonical MSC markers while excluding epithelial progenitors, enabling robust identification of this population (Figure 1B-C).

**Figure 1.**
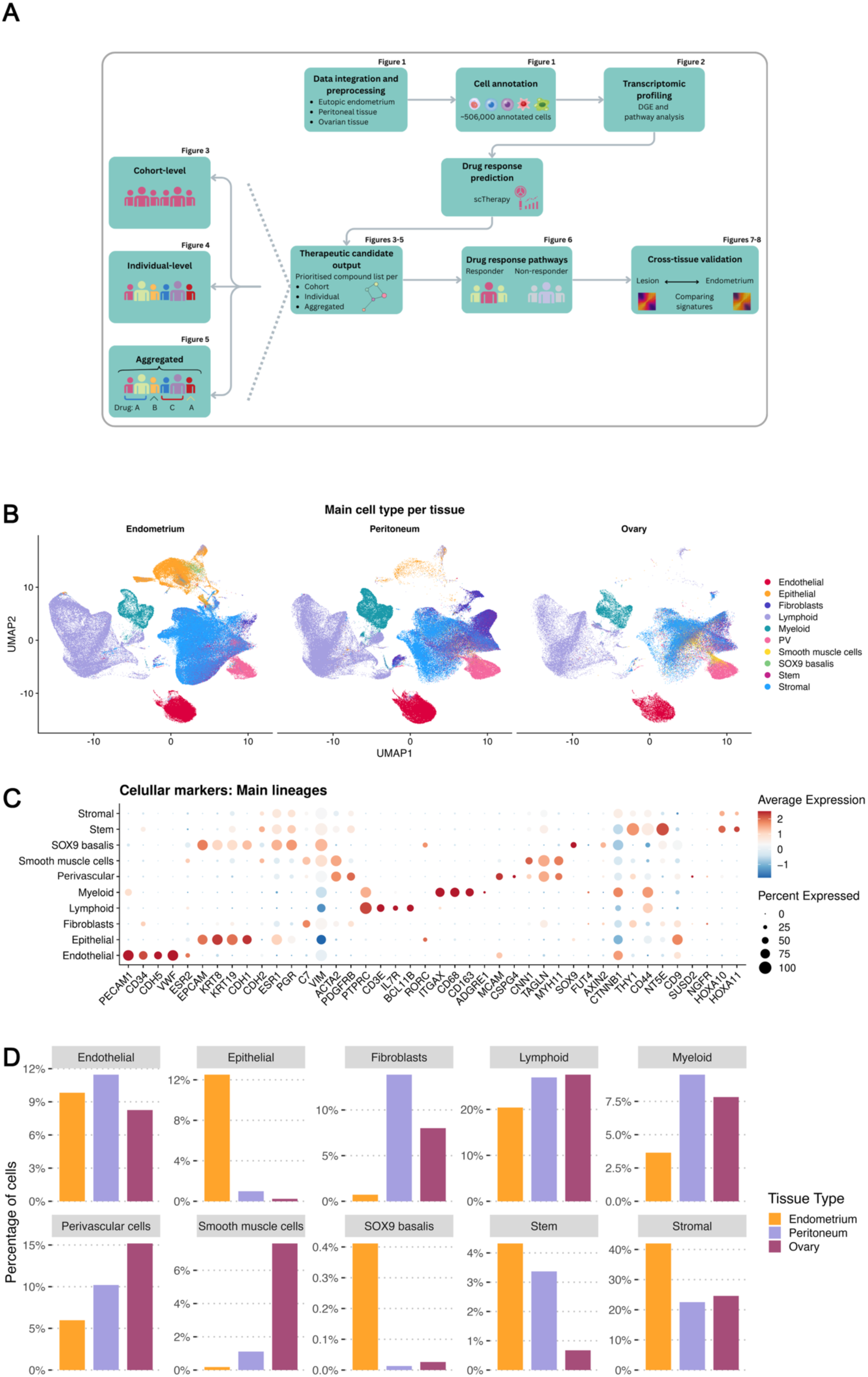
Integrated analysis workflow and cell-type resolution. **A)** Schematic overview of the analysis workflow. Corresponding figures are shown on the top left corner of each block. **B)** UMAP projections of the integrated single-cell dataset for each anatomical site, coloured by main cell type. The plots demonstrate clear separation of predicted cell types within each tissue, consistent with biologically meaningful clustering. Cell populations derived from HECA annotations were harmonised into broader categories, revealing distinct transcriptional profiles across stromal, epithelial, endothelial, immune and other compartments. **C)** Dot plot displaying the main cellular markers detected in the different cell lineages present in the scRNA-seq dataset. **D)** Percentage of main cell types per tissue type. PV: Perivascular cells, SOX9 basalis: SOX9+ Epithelial cells from the basalis part.

The annotation of the different cell types and their markers is displayed in Figure 1B, C. Cell lineages were annotated using canonical marker genes that capture the major compartments of endometrial tissue and lesions. Endothelial populations were defined by *PECAM1*, *CD34*, *CDH5*, *VWF*, and *ESR2*, while epithelial cells were identified through *EPCAM*, keratins (*KRT8*, *KRT19*), and hormone receptors (*ESR1*, *PGR*). Fibroblasts present in the basalis part of the endometrium, and the other tissues were distinguished by *C7*, *VIM*, *ACTA2*, and PDGFRB, with additional subsets including perivascular cells (*MCAM*, *PDGFRB*, *ACTA2*, *CSPG4*) and smooth muscle cells (*ACTA2*, *CNN1*, *TAGLN*, *MYH11*). Lymphoid cells *(PTPRC, CD3E, IL7R, BCL11B, RORC*) and myeloid populations (*ITGAX, CD68, CD163, ADGRE1*) captured the immune microenvironment, complemented by specialized subsets such as SOX9+ basalis cells (*SOX9, FUT4, AXIN2, CTNNB1*), stromal cells (*CD9, MCAM, SUSD2, NGFR, HOXA10, HOXA11*), and eMSCs (*THY1, CD44, NT5E*).

Overall, stromal cells represent the largest population across tissues, particularly in the eutopic endometrium, where they constitute approximately 40% of cells (Figure 1D). Endothelial and epithelial cells are also prominent, showing comparable proportions (10-12%) in endometrium, though epithelial cells are rare in peritoneal and ovarian tissues. Fibroblasts are more abundant in peritoneal tissues compared to the endometrium and the ovary. Lymphoid and myeloid lineage immune populations are enriched in peritoneal and ovarian tissues, reflecting immune cell infiltration. Perivascular and smooth muscle cells are less abundant populations across all tissues. SOX9+ basal and stem cells are present at low frequencies as expected, particularly in endometrium and peritoneum, highlighting the presence of progenitor compartments.

### Stromal, stem, and endothelial cells display disease-specific functional programmes linked to ECM remodelling and angiogenesis

Prior to assessing drug response, we characterised the transcriptional landscape of endometriosis to identify disease-associated gene expression patterns that could represent therapeutic targets. DGE analysis of pseudobulk single-cell profiles revealed that endometriosis-associated cells show distinct functional signatures compared to control subjects without endometriosis, across all major lineages (Figure 2). Epithelial cells showed upregulation of hormone-responsive and proliferative pathways, at higher rate than their typical role in cyclical remodelling and glandular activity (Hewitt *et al*., 2022; Mortlock, McKinnon and Montgomery, 2022). Immune populations, myeloid in particular, showed enrichment of immune and inflammatory response pathways, namely two previously reported in endometriosis; IL-6 and interferon signalling. Importantly, striking disease-associated changes were observed in the stromal and stem cell compartments. These populations displayed strong enrichment for epithelial-mesenchymal transition (EMT) and hypoxia pathways (Figure 2, Table 1), in line with the invasive and fibrotic features of endometriotic lesions previously reported (Chen *et al*., 2024; Vissers *et al*., 2024; Hosseinirad, Jeong and Barrier, 2025). We also observed lesion-associated changes in the endothelial population, including angiogenic signalling and vascular remodelling, reflecting the well-established importance of neovascularisation in lesion survival. These findings identify the stromal, stem and endothelial populations as principal therapeutic targets, while placing epithelial alterations in the broader context of hormone responsiveness. This focus is reinforced by our computational predictions and by existing evidence that stromal and vascular remodelling are critical for lesion establishment, fibrosis, and long-term persistence, thereby underscoring their translational significance.

**Figure 2.**
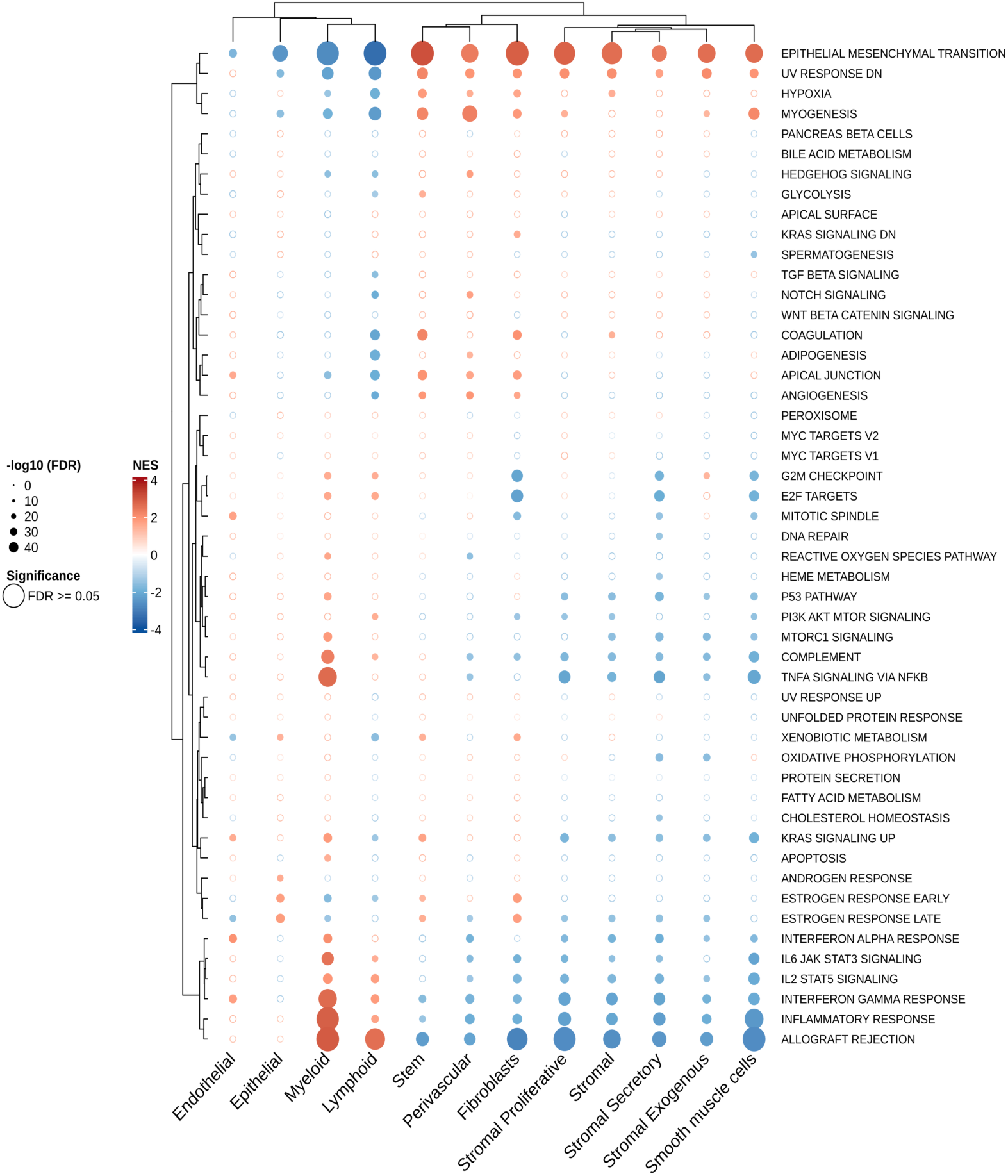
Pathway enrichment in endometriosis patients across cell types based on pre-ranked GSEA. Bubble heatmap showing enrichment of Hallmark pathways in major endometrial cell types. Rows represent the 50 gene signatures in the hallmarks collection; columns represent annotated cell types. Bubble colour reflects the normalised enrichment score (NES), while bubble size corresponds to the FDR-adjusted *p*-value. Empty bubbles indicate non-significant enrichment (FDR ≥ 0.05). Both rows and columns were hierarchically clustered using Euclidean distance on NES values to reveal shared functional programmes across cell types.

**Table 1.** Selected enriched pathways in cell types prioritised for drug prediction. Summary of prominent pathways identified through GSEA in endothelial, stromal and stem cell populations. These cell types were selected for downstream drug response prediction.

| *Main cell type* | *Enriched pathway* |
| --- | --- |
| **Endothelial** | Angiogenesis, TGFβ, Interferon, Inflammation. |
| **Stromal** | EMT, Hypoxia. |
| **Stem** | Angiogenesis, TGFβ, Hypoxia, Glycolysis, Oestrogen response, Xenobiotic metabolism, KRAS signalling. |

### Cohort-level single-cell drug response prediction reveals broad and tissue-specific therapeutic candidates

Based on their central roles in endometriosis pathophysiology and their representation across all tissues, subsequent analyses focused on the stromal, stem and endothelial cell populations. To identify compounds capable of reversing disease-associated transcriptional states within these key populations, we applied scTherapy using DGE profiles derived from endometriosis versus control samples, stratified by tissue (Figure 1A). We also filtered out drug–cell combinations predicted to be toxic, retaining only predictions in which a given cell type within a patient showed a high response probability without associated toxicity. This enabled cohort-level prioritisation of candidate therapeutics across individuals, cell types, and tissue sites.

Across the cohort, the most frequently prioritised compound was ABT-751, predicted in approximately 85% of patient samples (Figure 3A). Panobinostat and JBJ-26481585 followed next, with approximately 25% predicted efficacy. Other drugs, including romidepsin, delanzomib, and bortezomib, appeared less frequently, indicating narrower predicted efficacy across patients. When predictions were stratified by cell type (Figure 3B), ABT-751 displayed broad activity in all three cell types, with strongest efficacy in endothelial cells. Panobinostat showed high predicted efficacy in stromal and stem cells, while JNJ-26481585 had a balanced effect across populations. In contrast, agents such as docetaxel and SN-38 exhibited more restricted, cell-type-specific profiles.

**Figure 3.**
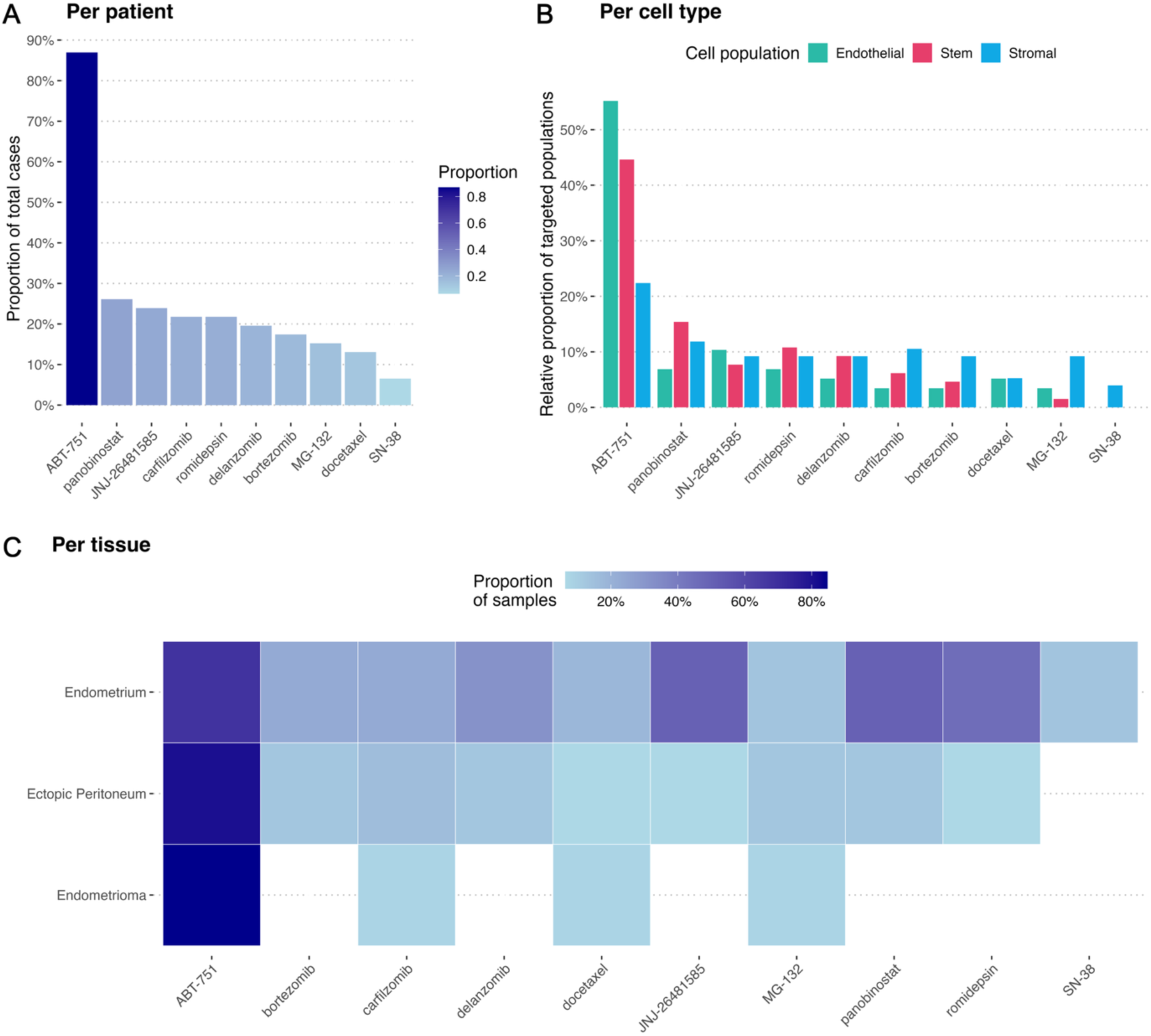
Summary of personalised drug predictions across all patients, cell populations and tissue types. **A)** Bar plot showing the proportion of patients for which each compound is predicted to be effective in at least one cell population (endothelial, stem, and/or stromal). **B)** Relative proportion of targeted cell populations (stromal, endothelial, or stem) for each compound. **C)** Proportion of samples across all individuals that contain at least one cell type predicted to respond to each compound.

Drug response patterns further revealed tissue-dependent differences (Figure 3C). Eutopic endometrium exhibited sensitivity to a broader range of compounds. Ectopic peritoneal cells were particularly responsive to ABT-751 and to lesser extent to other drugs such as panobinostat and carfilzomib, while ovarian lesions showed a more selective response pattern dominated by ABT-751. These findings suggest that despite overarching therapeutic trends, drug response in endometriosis is strongly influenced by both cellular identity and tissue context, pointing to additional layers of heterogeneity at the individual level. To explore this inter-patient variation in greater detail, we next applied scTherapy on a per-patient basis to characterise individual therapeutic sensitivity profiles.

### Tissue context and individual variation drive differential drug responses

Because therapeutic responses vary across individuals and tissues, we investigated inter-individual differences in therapeutic sensitivity by applying scTherapy at the patient level across tissue types and cellular compartments (Figure 4). Our results indicate distinct patterns of drug sensitivity between patients. In eutopic endometrium (Figure 4A) the leading candidates were romidepsin, JNJ-26481585, and panobinostat, all HDAC inhibitors. A smaller subset of patients responded preferentially to ABT-751 or proteasome inhibitors such as bortezomib and delanzomib. By contrast, in ectopic peritoneal lesions (Figure 4B), responses were more evenly distributed across a narrower set of compounds with ABT-751, bortezomib, and panobinostat dominating the top predictions. This pattern indicates a more convergent therapeutic landscape in ectopic tissue compared to eutopic endometrium. Notably, excepting the flagged drugs due to predicted toxicity in healthy tissue (romidepsin, JNJ-26481585, and bortezomib), panobinostat emerged as a recurrent candidate across both ectopic and eutopic compartments. In the ectopic ovary (Figure 4C), a different sensitivity profile was observed with bortezomib, delanzomib and carfilzomib emerging as the most frequently predicted effective agents. Notably, ABT-751 and panobinostat, which were consistently prioritised in other tissues, appeared in a smaller proportion of ovarian samples, suggesting reduced susceptibility of ovarian-derived cells to microtubule and HDAC inhibition.

**Figure 4.**
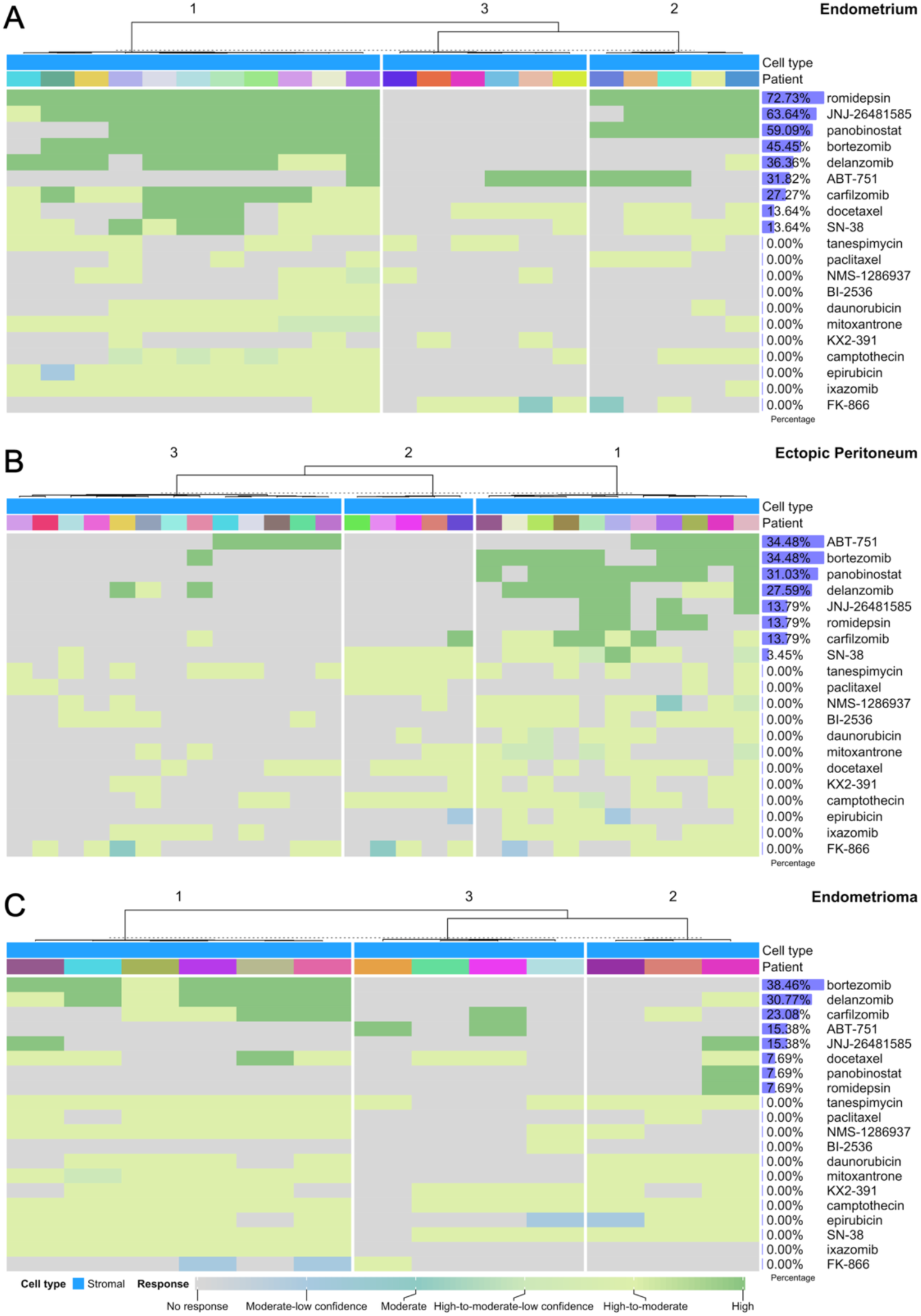
Patient-stratified profiles of predicted drug responses. Heatmaps depicting responses in stromal cells of **A)** eutopic endometrium, **B)** ectopic peritoneal tissue, and **C)** ectopic ovarian tissue. Each heatmap displays the top 20 drugs selected for that specific tissue type. Drugs were filtered to include those with a response variability (Shannon’s entropy) ≥ 0.5 and then ranked by the total number of high or high-to-moderate responses. Columns represent individual patient samples. Columns are clustered into three k-means groups based on response profile similarity. Percentage denotes percentage of patients that exhibited a high response for each drug.

Overall, these results demonstrate that predicted drug efficacy varies both across tissues and between individuals, reflecting underlying differences in cellular composition, local microenvironment, and molecular state. The heterogeneity observed here underscores the limitation of a uniform therapeutic strategy for endometriosis.

### Clustering of the personalised predictions reveal recurrent candidates

Having established marked inter-individual variation in predicted drug sensitivities across tissues, we next sought to determine whether these heterogeneous responses nevertheless converged into a limited number of recurring therapeutic patterns. Specifically, we asked whether patients could be grouped according to shared drug-response profiles within each major cell population, independent of tissue origin (Figure 1A). This approach allowed us to assess whether stem, endothelial, and stromal cell populations exhibit conserved molecular vulnerabilities across patients, even when derived from distinct tissue microenvironments.

For each cell population, we selected the top 10 compounds predicted to elicit the highest therapeutic response without associated toxicity and performed unsupervised clustering of the individual-level prediction matrices (Figure 5). This analysis identified distinct patient clusters enriched for overlapping candidate drugs, suggesting that therapeutic heterogeneity may reflect a small number of conserved response programmes rather than purely random variability. Notably, HDAC inhibitors (e.g. panobinostat, romidepsin) and tubulin polymerisation inhibitor ABT-751 consistently co-occurred across clusters, implying potential convergence on shared mechanistic pathways or complementary targets. In the case of stem cells (Figure 5A), response profile in Cluster 1 was dominated by ABT-751 whereas Cluster 2 was characterised by a more diverse sensitivity profile, with a notable response to panobinostat. Endothelial cells (Figure 5B) showed high response to JNJ-26481585 and SN-38 in Cluster 1 and highest response to ABT-751 in Cluster 2, albeit a less dominant response than in stem cells. On the other hand, stromal cells (Figure 5C) exhibited multi-compound sensitivity in Cluster 1, and MG-132-dominant response in Cluster 2. These findings support the existence of recurring, cell-type-specific therapeutic profiles in endometriosis that transcend tissue context, providing a rational basis for stratified or combinatorial treatment approaches.

**Figure 5.**
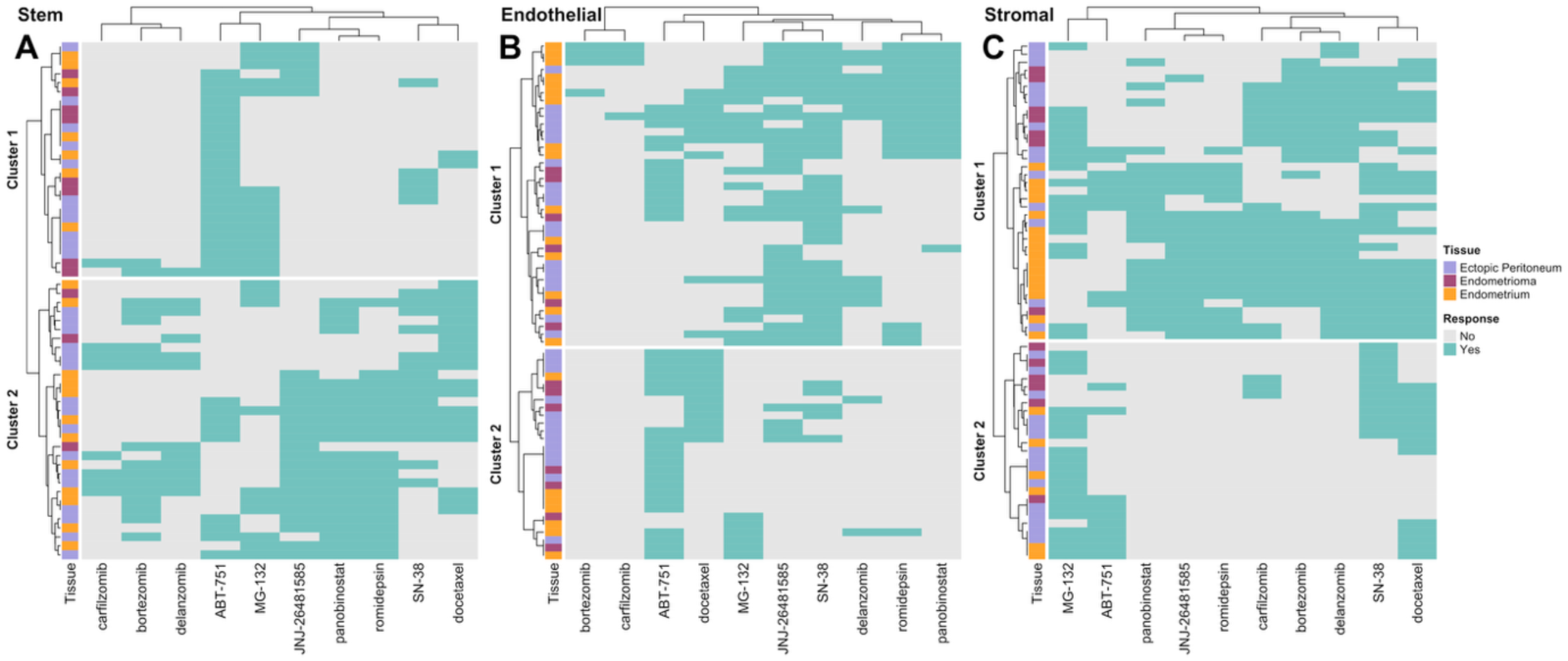
Hierarchical clustering of drug response profiles. Binary heatmaps illustrating the drug response patterns for **A)** stem, **B)** endothelial and **C)** stromal cells. Each row represents a patient sample from a given anatomical site of origin, and each column represents a unique high-effect, non-toxic drug. Teal represents a positive drug response; gray represents no response. Samples (rows) are grouped into two primary clusters, as predefined, based on response similarity using hierarchical clustering.

### Molecular features underlying response to ABT-751 and panobinostat were driven by proliferation and fibrosis

To validate the predicted efficacy of the top-performing drug candidates, we focused on ABT-751 and panobinostat. ABT-751 was inferred to exert therapeutic effects across multiple cell types and patient groups, showing consistent prioritization across diverse tissues. In contrast, panobinostat emerged as a recurrent candidate in both ectopic and eutopic compartments. Together, these two compounds were predicted to effectively target approximately 51.6% of stromal cells derived from peritoneal lesions. We focused on these lesions because they provided a sufficient number of samples to ensure statistically robust downstream analyses, in contrast to the ovarian lesions. We concentrated on stromal cells as they constitute the predominant cell population within these lesions. To further elucidate the molecular basis of inter-patient heterogeneity, we compared the transcriptional profiles of predicted responders and non-responders to ABT-751 and panobinostat. This approach aimed to identify biomarkers and mechanisms underlying the differential drug sensitivities observed in our predictive analyses.

Stromal cells from peritoneal lesions of patients predicted to respond to ABT-751 exhibited a transcriptional program characterised by enhanced ECM remodelling, inflammation and fibrotic activation. Upregulated genes included *POSTN* (Logan, Yango and Tran, 2017; She *et al*., 2024), key mediators of collagen organisation and tissue fibrosis, alongside *DKK1* (Lee *et al*., 2025), *TSPAN13* (Zhang *et al*., 2024), and lncRNA *H19* (Wen *et al*., 2024), which promote inflammation, proliferation and metabolic programming (Figure 6A). In contrast, downregulation of genes associated with immune regulation and apoptotic clearance, including *SPINT2* (Roversi *et al*., 2013), *BEX2* (Tamai *et al*., 2020; Fukushi *et al*., 2021), *GATA4* (Babu *et al*., 2025), *STAR* (Zhong, Ding and Xia, 2025), *CHRM3* (Wang *et al*., 7 2015), and *PLA2G5* (Sato *et al*., 2014), indicated disrupted protease balance, steroidogenesis, and immune surveillance. Over-representation analysis of upregulated genes in high-responder stromal cells confirmed significant enrichment in pathways related to collagen metabolism, ECM organisation, and vascular development (Figure 6B). *CytoTRACE2*, a computational framework for estimating cellular stemness based on transcriptomic profiles, was used to quantify the differentiation potential of the cells. The analysis revealed a mixed population of oligopotent and differentiating stromal cells suggesting that ABT-751 responsive lesions maintain proliferative and regenerative potential (Figure 6C), as evidenced by the presence of stromal cells at different developmental stages. In contrast, stromal cells in the non-responder group appear to remain in a multipotent state, without showing populations at more advanced stages of differentiation. Thus, the stromal cells responding to ABT-751 likely have a pro-inflammatory and proliferative phenotype.

**Figure 6.**
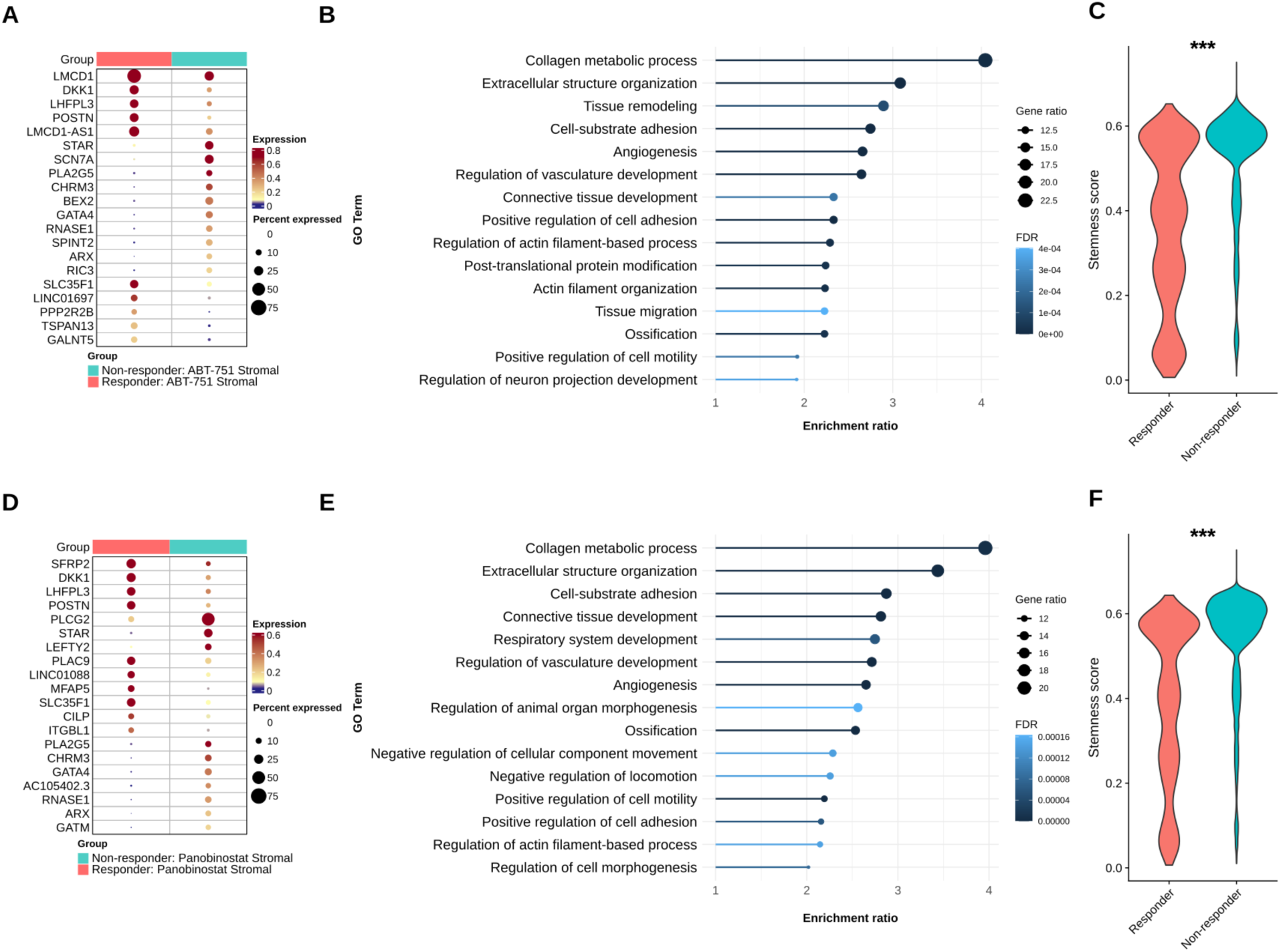
Transcriptomic profiles of peritoneal lesion-derived stromal cells from inferred high and non-responders to ABT-751 and panobinostat. Heatmap showing the top 10 up- and downregulated genes between high and non-responder groups to **A)** ABT-751 and **D)** panobinostat. Enrichment analysis of upregulated genes in high responders to **B)** ABT-751 and **E)** panobinostat. Stemness score of stromal cells calculated by *CytoTRACE2* for **C)** ABT-751 response (two-sided Kolmogorov–Smirnov test, *p* < 2 × 10⁻¹⁶; Cliff’s d = –0.51) and **F)** panobinostat response (two-sided Kolmogorov–Smirnov test, *p* < 2 × 10⁻¹⁶; Cliff’s d = –0.52).

Similarly, stromal cells of peritoneal lesions from inferred responders to panobinostat displayed coordinated upregulation of genes involved in ECM organisation, proliferation and angiogenesis. *DKK1* and *POSTN* were consistently overexpressed, mirroring their elevation in ABT-751 responders, and reinforcing a shared pro-fibrotic phenotype (Figure 6D). Additional upregulated genes included *SFRP2* (Heinosalo *et al*., 2018), linked to WNT5A-mediated lesion growth, and *CILP* (Liu *et al*., 2021), associated with matrix deposition and fibrotic remodelling. Conversely, *GATA4* (Babu *et al*., 2025), a positive regulator of mesenchymal stem cell senescence was downregulated in the responder group, suggesting cells remain in a quiescent state that retain the capacity to differentiate. The opposite phenomenon is likely observed in non-responders in which the cells have lost their capacity to differentiate. Pathway enrichment analysis confirmed activation of collagen metabolism; hallmarks associated with tissue migration, such as, positive regulation of cell motility and angiogenic processes (Figure 6E), while *CytoTRACE2* analysis indicated a population spanning oligopotent to differentiated states (Figure 6F). These results suggest that stromal cells that are predicted to respond to panobinostat have a proliferative, pro-fibrotic transcriptome resembling active development transcriptional.

### Panobinostat and ABT-751 response is conserved in stromal cells located in the eutopic endometrium and ectopic peritoneal lesions

To evaluate whether eutopic endometrium can serve as a non-invasive source for assessing drug response, we tested whether the transcriptomic changes induced by the two in-silico–predicted drugs in peritoneal lesion-derived stromal cells could also be detected in eutopic endometrial stromal cells from endometriosis patients. To address this, we first compared their pseudobulk transcriptomic profiles, stratified by menstrual cycle phase. Pearson correlation analysis revealed a higher similarity between stromal cells from peritoneal lesions and eutopic endometrium in the secretory phase, whereas proliferative-phase cells showed lower correlation (Figure 7A). Furthermore, key stromal and cycle-phase markers were conserved across both tissues (Supplementary Figure 2). Overall, the transcriptomic signature of endometrial stromal cells closely mirrors that of lesion-derived stromal cells, suggesting that endometrial stromal cells could serve as a non-invasive surrogate for drug testing.

**Figure 7.**
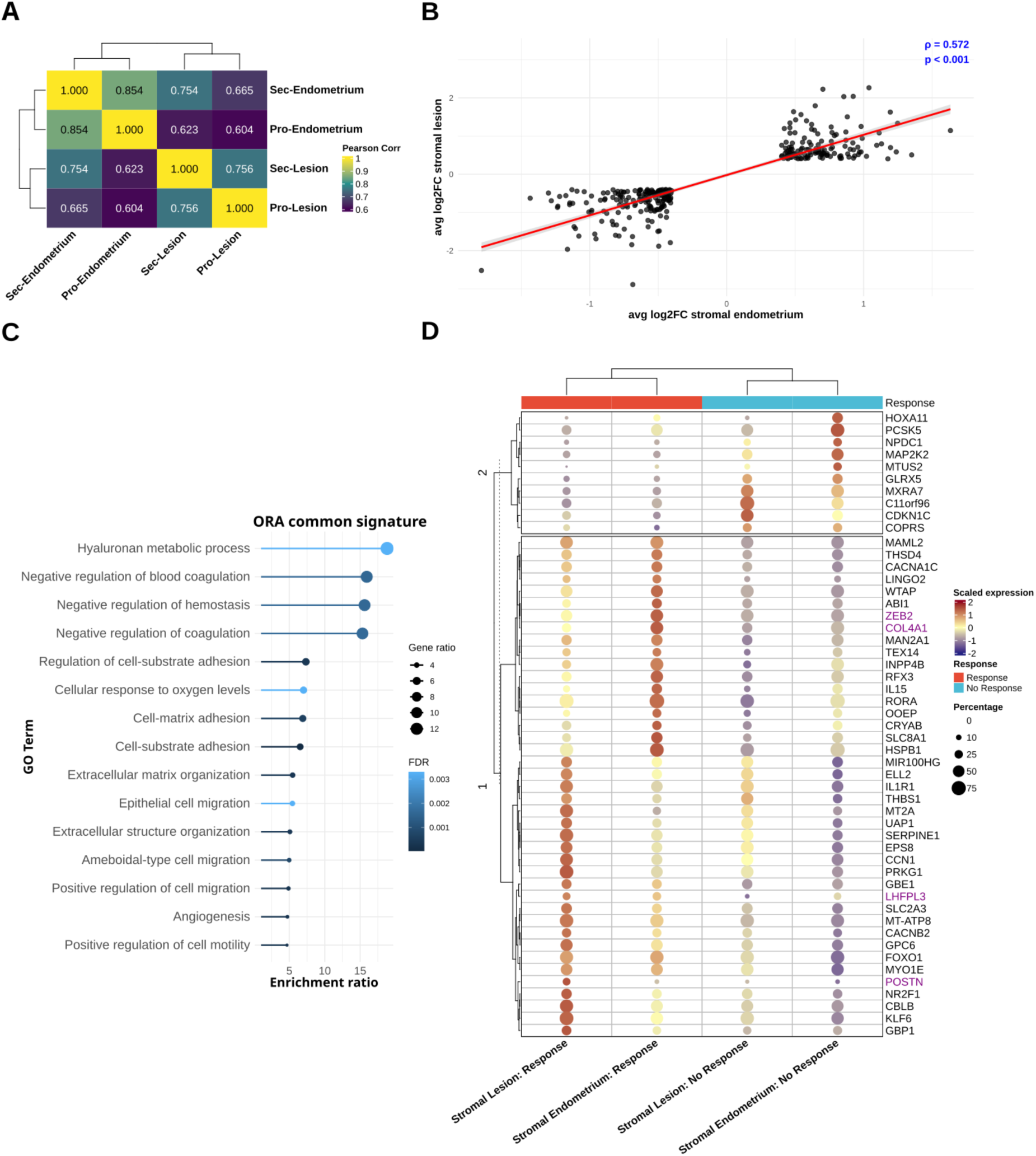
Conserved transcriptional response to panobinostat across stromal cells from eutopic and ectopic peritoneal tissues. **A)** Pearson correlation of the top 4,000 most variable genes expressed in stromal cells from peritoneal lesions and eutopic endometrium across different menstrual cycle phases (secretory and proliferative phases). **B)** Correlation of differential expression results for the intersecting genes identified as DEGs in stromal cells from peritoneal lesions and eutopic endometrium (Kendall correlation, τ = 0.57177, *p* < 2.2 × 10⁻¹⁶). **C)** ORA of genes upregulated in stromal cells from high-response patients to panobinostat, overlapping with genes expressed in peritoneal lesions and eutopic endometrium. **D)** Heatmap showing the top 50 most significant DEGs shared between stromal cells in both tissues. Genes highlighted in bold purple indicate top DEGs in peritoneal lesions or genes previously associated with endometriosis. **Sec-endometrium**: stromal cells located in the endometrium in secretory stage. **Pro-Endometrium**: stromal cells located in the endometrium in proliferative stage. **Sec-Lesion**: stromal cells located in endometriosis lesion in the peritoneum in secretory stage. **Pro-Lesion**: stromal cells located in endometriosis lesion in the peritoneum in proliferative stage.

Comparative DGE analysis between high -and non-responders identified substantial overlap in the transcriptional response to panobinostat across the two tissues. Among upregulated genes, 135 followed the same direction of change (hypergeometric test *p* < 8.68x10^-8^, Jaccard index = 0.15), while 184 downregulated genes showed consistent trends (hypergeometric test *p* < 9.04x10^-3^, Jaccard index = 0.16). Correlation of shared genes yielded a significant positive association (Figure 7B). Panobinostat induced robust and consistent changes in gene expression across tissues, indicating a non-random, biologically coordinated molecular response.

ORA of the up-regulated shared genes between the two tissues revealed enrichment for biological processes related to angiogenesis, ECM organisation, and cell migration, pathways closely linked to lesion establishment and fibrotic remodelling (Figure 7C). Moreover, clustering of the transcriptomic signatures grouped samples primarily by responder status rather than tissue origin (Figure 7D). Genes such as *POSTN*, *COL4A1*, and *ZEB2* were consistently upregulated in high responders from both tissues, supporting a shared pro-fibrotic and invasive phenotype.

Finally, we compared the transcriptomic similarity between responders and non-responders to ABT-751 within stromal cells from endometriosis patients, using the same analytical strategy described in the previous section. DEGs were identified for responders versus non-responders among endometrial stromal cells and compared to the transcriptomic signature obtained from peritoneal lesion stromal cells. We observed a substantial overlap in the transcriptional response to ABT-751 across both tissues. Among upregulated genes, 230 showed concordant direction of change (hypergeometric test, *p* = 4.43 × 10⁻³³; Jaccard index = 0.252). Notably, 381 downregulated genes displayed a highly significant overlap across tissues (hypergeometric test, *p* = 8.70 × 10⁻⁴⁰), with a moderate degree of similarity (Jaccard index = 0.303). Correlation analysis of shared genes yielded a significant and high correlation (Figure 8A). These findings indicate that ABT-751 induces robust and coordinated gene expression changes across tissues, supporting a biologically consistent molecular response.

**Figure 8.**
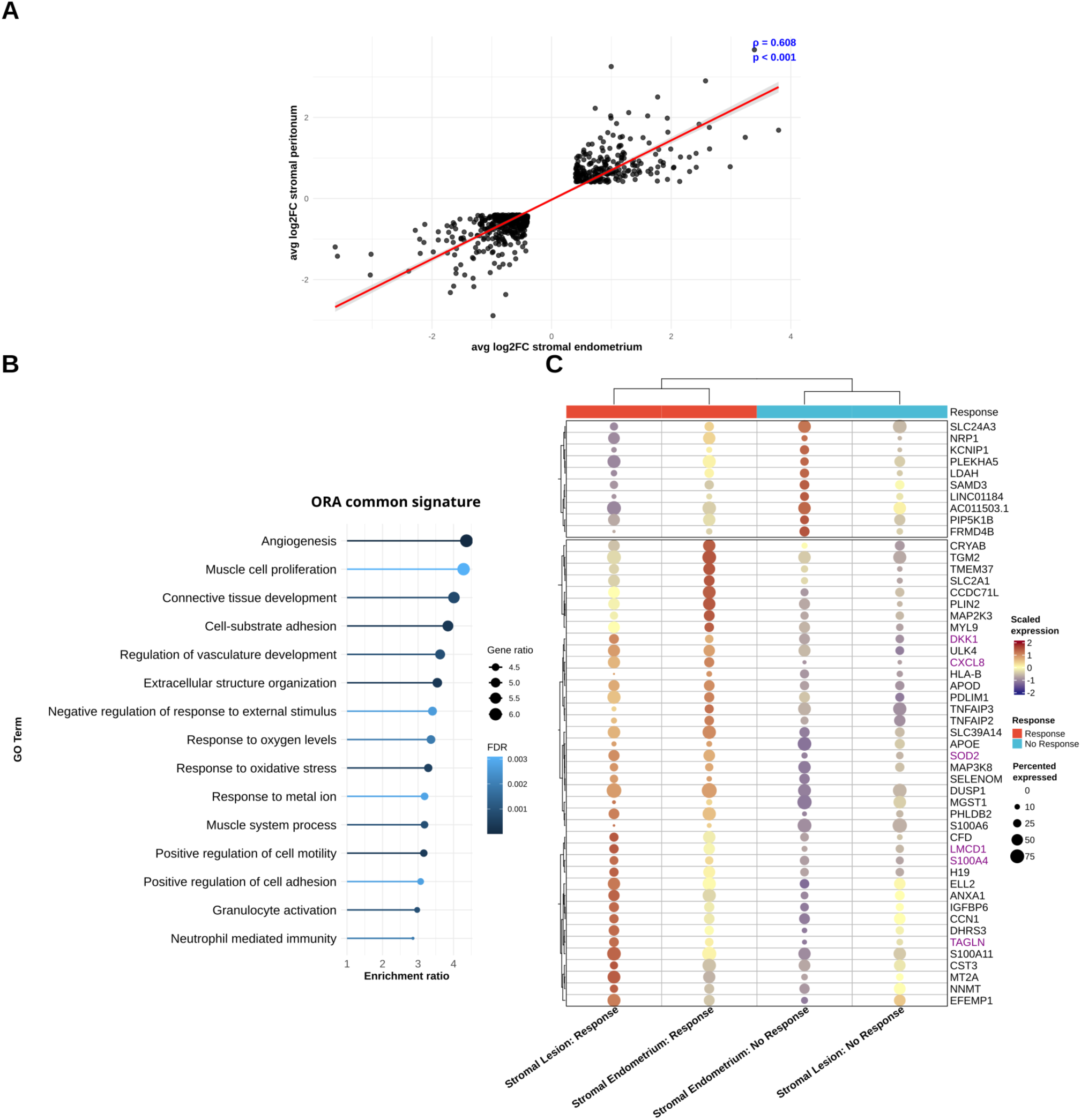
Conserved transcriptional response to ABT-751 across stromal cells from eutopic and ectopic peritoneal tissues. **A)** Correlation of differential expression profiles between stromal cells from peritoneal lesions and eutopic endometrium in ABT-751 responders (Kendall correlation, τ = 0.607, *p* < 2.2 × 10⁻¹⁶). **B)** ORA of upregulated genes shared between eutopic and ectopic stromal cells from high-response patients, highlighting enrichment in angiogenesis, ECM organisation, and cell migration pathways. **C)** Heatmap showing the top 50 most significant DEGs shared between stromal cells in both tissues. Genes highlighted in purple denote top-ranked DEGs in peritoneal lesions or those previously indicated in endometriosis.

Enrichment analysis of upregulated genes shared between endometrial and lesion-derived stromal cells revealed significant enrichment for biological processes related to angiogenesis, extracellular matrix organization, and increased cell mobility and adhesion (Figure 8B). These findings align with the functional patterns observed in the shared gene signature. Among the overlapping genes, several previously implicated in endometriosis and fibrotic processes were identified, including *DKK1* and *LMCD1*, as well as *CXCL8* (Bordon, 2023) whose inhibition has been shown to attenuate the fibrotic environment characteristic of endometriosis. Moreover, *SOD2* (superoxide dismutase 2), a key mitochondrial enzyme for the detoxification of reactive oxygen species that promotes angiogenesis, was upregulated in stromal cells from ABT-751 responders (Connor *et al*., 2005). Finally, *TAGLN* (Transgelin), a gene associated with cellular invasion and migration, was also upregulated in responder stromal cells, consistent with its previously reported involvement in endometriosis progression (Muraoka *et al*., 2023).

These results suggest that the core transcriptomic signature of the drug response can be recapitulated in stromal cells from the eutopic endometrium of endometriosis patients, supporting their use as a minimally invasive source for evaluating candidate therapeutics for the disease.

## Discussion

In this study, we demonstrate the feasibility and translational potential of integrating patient-derived single-cell transcriptomes with drug-induced transcriptional signatures to systematically investigate therapeutic heterogeneity in endometriosis. We aimed to determine whether variability in predicted drug responses reflects underlying molecular differences across patients and lesions. Our findings provide evidence that endometriosis is characterised by pronounced inter-patient and intra-lesion heterogeneity at the transcriptional level, and that this heterogeneity translates into distinct therapeutic vulnerabilities. By investigating drug response predictions at single-cell resolution, we uncover the molecular programmes that shape sensitivity to specific compounds and begin to outline the molecular basis for the variable treatment outcomes in this disease.

Approved treatments for endometriosis remain largely limited to hormonal modulation, with synthetic progestins such as Dienogest as first-line therapy and gonadotrophin-releasing hormone (GnRH) agonists such as Leuprolide acetate and Goserelin acetate as additional options. Newer combination therapies, including the GnRH antagonist Relugolix with add-back therapy, offer symptom relief but are not curative, require prolonged administration, and are associated with significant side effects that impact fertility and bone health (Sauerbrun-Cutler and Alvero, 2019). Recurrence rates remain high after treatment cessation. Despite decades of research, the therapeutic pipeline for endometriosis remains heavily dominated by hormonal agents and recent non-hormonal candidates failed in clinical trials (Goenka, George and Sen, 2017; Malvezzi *et al*., 2020). For example, HSD17B1 inhibitors, selective progesterone receptor modulators (SERMs), and aromatase inhibitors have yielded inconsistent benefits, reflecting the limitations of targeting single pathways in a disease marked by heterogeneity and variable patient responses. Current clinical care and drug development pipelines do not incorporate molecular profiling, yet our results show that endometriosis comprises several transcriptionally distinct patient groups. Defining and stratifying these groups will be essential for matching patients to the most appropriate mechanism of action, mirroring the principles that have transformed precision oncology. In this context, our analyses identified non-hormonal compounds, such as HDAC inhibitors, proteasome inhibitors and tubulin polymerisation inhibitors, with wide predicted vulnerability up to approximately 45% of stromal and stem cell populations. These compounds not only exhibit mechanistically distinct activity but also tissue-specificity, representing promising directions for targeted, non-hormonal interventions.

A key feature of our approach is that drug responses are predicted at the level of individual cell types. This is particularly important in endometriosis, where the cellular composition is highly heterogeneous, dynamic and shaped by tissue-specific microenvironments. Therefore, therapeutic effects may differ between the different cellular compartments and anatomical sites. Another important consideration for therapeutic development is that endometriosis is not confined to a single anatomical site. Lesions arise in the peritoneum, ovary, and deep pelvic tissues, or extra-pelvic regions, while the eutopic endometrium itself also displays aberrant transcriptional and functional features. Our analyses revealed that predicted drug sensitivities varied not only across cell types but also between lesion locations and the eutopic tissue. For example, panobinostat displayed selective efficacy in peritoneal lesions and eutopic endometrium but limited effects in the ovarian lesions, whereas ABT-751 targeted all three tissues. Moreover, eutopic tissue was more sensitive to HDAC inhibitor class drugs. This highlights the necessity of designing treatment strategies that can address both ectopic and eutopic tissues. Given that the eutopic endometrium contributes cells that seed new lesions and fuel symptoms, while established ectopic lesions drive pain and infertility through inflammation, fibrosis and local invasion, a comprehensive treatment must target both. In this context, ABT-751 and panobinostat proved as mechanistically-driven compounds with broad coverage at tissue and/or cell level to potentially tackle endometriosis more effectively.

Notably, many of the compounds identified, including ABT-751 (Mahgoub *et al*., 2024), bortezomib (Aubrey *et al*., 2025), MG-132 (Marzan *et al*., 2025), and HDAC inhibitors like romidepsin (Sequera *et al*., 2025) and panobinostat (Demir *et al*., 2024), are either FDA-approved or have been previously explored in other clinical contexts. Furthermore, some of the compounds have been linked to endometriosis previously, validating the robustness of the *in silico* pipeline. ABT-751 was reported as a potential drug against endometriosis in a 2022 study by Wang and colleagues (Wang *et al*., 2022). In an endometriosis rat model, bortezomib was reported to reduce lesion growth (Celik *et al*., 2008). More recently, a computational drug repurposing pipeline based on protein-protein interactions identified bortezomib as one of the compounds inferred as a candidate for a number of diseases, including endometriosis (Ortaakarsu and Medetalibeyoğlu, 2024). MG-132 was previously assessed for its role in modulating inflammation and autophagy in endometriotic stromal cells (Z.-X. Huang *et al*., 2023).

While panobinostat was considered in high-throughput screening assays, it has not been explored yet as a therapeutic agent for endometriosis (Churchill *et al*., 2023). Several lines of evidence support the relevance of HDACs in endometriosis. For example, HDAC1 was reported to be markedly overexpressed in endometriotic lesions, particularly in epithelial and stromal cells, whereas HDAC2 showed more variable results (Colón-Díaz *et al*., 2012). Previous research on prostate cancer cells demonstrated that exposure to HDAC inhibitors resulted in demethylation of promoters investigated in the study (Sarkar *et al*., 2011). These findings provide a strong translational rationale for the use of HDAC inhibitors, especially those with HDAC1 selectivity, as potential therapeutic agents in endometriosis. Elevated HDAC1 is thought to promote pathological chromatin repression and gene silencing, and its inhibition offers an opportunity to restore normal gene expression (Lawlor and Yang, 2019). Clinical observations in adenomyosis further support this mode of action. Valproic acid, a potent HDAC inhibitor, was shown to reduce uterine size, menstrual volume and dysmenorrhea severity in patients (Liu, Yuan and Guo, 2010). Importantly, the relevance of epigenetic-targeting drugs is further reinforced by our recent findings in menstrual blood-derived stem cells, which demonstrated hypermethylation affecting pathways linked to proliferation, invasion and aberrant cell-state maintenance (Tiniakou *et al*., 2025). This provides additional support for therapies that act on epigenetic dysregulation. Thus, the use of HDAC inhibitors may counteract both histone- and DNA-methylation mediated repression, offering a promising strategy to restore regulatory balance in endometriosis. Follow-up *in vitro* and *in vivo* work will be critical to assess the feasibility and therapeutic potential of these compounds in treating endometriosis.

Beyond predicted efficacy, we aimed to explore the transcriptomic differences between responder and non-responder patients to identify genes associated with drug sensitivity. Given the in-silico nature of our analysis, we focused on peritoneal lesions and stromal cells for two main reasons. First, the peritoneal lesion cohort included enough patients (n = 29) to enable statistically robust comparisons. Second, stromal cells constitute the most abundant cell type across endometrial compartments, representing a biologically relevant population for assessing transcriptional determinants of drug response. Finally, we focused our analysis on ABT-751 and panobinostat, as their patient-stratified response profiles suggested that the combined use of these two compounds could effectively target approximately 51.7% of patients with peritoneal endometrial lesions. The transcriptomic profiling of inferred responders versus non-responders provided mechanistic insight into the cellular states associated with drug sensitivity.

Our comparative transcriptional analysis of predicted responders and non-responders revealed distinct molecular signatures that may explain inter-patient heterogeneity in therapeutic response to ABT-751 and panobinostat. Stromal cells from peritoneal lesions predicted to respond to both compounds exhibited convergent transcriptional programs characterized by enhanced ECM remodelling, fibrotic activation, and proliferative capacity. The consistent upregulation of key fibrotic mediators such as *POSTN* and *DKK1* across both responder groups suggests that a pro-fibrotic, actively remodelling stromal phenotype may be a critical determinant of drug sensitivity in endometriotic lesions. Interestingly, this dysregulation of *POSTN* and *DKK1* aligns with current evidence demonstrating that *POSTN* dysregulation is linked to EMT in endometriosis (Zheng *et al*., 2016) and promotes migration, invasion, and increased adhesion of stromal cells within peritoneal lesions (Xu *et al*., 2015), while *DKK1* is upregulated in the serum of women with endometriosis (Kasoha *et al*., 2019). Notably, *CytoTRACE2* analysis revealed that responsive stromal populations maintain a dynamic differentiation landscape, spanning oligopotent to differentiating states, which likely reflects ongoing proliferative and regenerative activity within these lesions. In contrast, non-responsive stromal cells appeared transcriptionally quiescent, exhibiting sustained multipotency without progression toward differentiation.

Finally, the transcriptomic signature associated with either ABT-751 or panobinostat responsiveness was recapitulated in stromal cells from the eutopic endometrium. The shared up- and downregulated gene sets were significantly correlated, suggesting a conserved molecular programme triggering a consistent therapeutic response irrespective of tissue context. This reinforces the concept that eutopic stromal cells retain pathological features reflective of ectopic lesions and can serve as a reliable, accessible model for preclinical testing of candidate therapeutics in endometriosis. Limiting the need for surgical samples and enabling the use of non-invasive samples such as menstrual blood, would facilitate mechanistic studies and transform translational research thereby accelerating drug screening for endometriosis.

Our study still has important limitations that need addressing. First, the computational predictions remain to be validated experimentally *in vitro* and *in vivo*. Second, the integrated dataset relies on the quality and consistency of public datasets, which may vary in terms of sequencing depth, cell capture efficiency, and completeness of clinical metadata. Third, the scope of the analysis is constrained by the compounds available in existing drug perturbation databases, meaning that potentially relevant therapeutics not captured in these resources could not be assessed. Finally, the perturbation profiles used to train scTherapy are derived largely from cancer cell lines. Although this may introduce a bias towards oncogenic signalling contexts, the relevance of these signatures is supported by the biological and therapeutic overlap between cancer and endometriosis. Future studies should prioritise functional validation, ideally using primary endometriosis models or organoid systems, and ultimately assess the efficacy and safety of prioritised candidates in preclinical biosimilar models or clinical trials.

In summary, our study provides a framework for dissecting therapeutic heterogeneity in endometriosis at single-cell resolution, revealing how patient-, tissue- and cell-type-specific transcriptional states shape drug susceptibility. By uncovering conserved stromal transcriptomic signals linked to predicted treatment response, our findings highlight the central role of stromal biology in determining therapeutic efficacy and support the use of eutopic endometrium as an accessible surrogate for assessing drug response. These results provide a rationale for integrating molecular profiling into future drug development pipelines, moving beyond the current one-size-fits-all paradigm and toward stratified, molecularly informed treatment strategies.

