## Supplementary material for "Beyond one-size-fits-all: single-cell transcriptomic signatures predict drug efficacy and reveal responder subgroups in endometriosis"

**Supplementary Table 1. Distribution of sample types and metadata across the datasets.**

| *Dataset ID* | *Menstrual cycle phase (n)* | *Endometriosis stage (n)* | *Contraceptives (n)* | *Sample type (n)* | | | | |
|---|---|---|---|---|---|---|---|---|
| | Proliferative/ Secretory/ Other | Stage I/II/III/IV | Yes/No | Ectopic adjacent | Ectopic endometrium | Ectopic ovary | Eutopic endometrium | Unaffected ovary |
| *GSE214411* | 6/7/0 | 5/1/0/0 | 0/13 | 0 | 0 | 0 | 13 | 0 |
| *GSE213216* | 22/16/11 | 3/11/13/16 | 11/38 | 4 | 23 | 8 | 10 | 4 |
| *GSE179640* | 9/6/16 | 0/3/3/22 | 27/4 | 7 | 8 | 4 | 12 | 0 |
| *PRJNA932195* | 0/0/17 | 0/0/4/12 | 1/16 | 0 | 13 | 4 | 0 | 0 |

**Supplementary Table 2. Performance metrics for SVM classification of HECA-defined cell types in the test set.** Precision, recall, and F1-score are reported for each of the 36 HECA annotation classes. The number of test samples (cells) assigned to each class is also shown. To summarise classifier performance, both macro and weighted averages are included: the macro average reflects the unweighted mean across all classes (suitable for balanced datasets), while the weighted average accounts for class imbalance by incorporating the number of test samples per class.

| *HECA annotation* | *Precision* | *Recall* | *F1-Score* | *Test samples* |
|---|---|---|---|---|
| ***Arterial*** | 0.93 | 0.92 | 0.93 | 119 |
| ***Ciliated*** | 0.97 | 0.97 | 0.97 | 118 |
| ***Cycling*** | 0.7 | 0.63 | 0.67 | 128 |
| ***Fibroblast_basalis*** | 0.97 | 0.93 | 0.95 | 118 |
| ***Glandular*** | 0.88 | 0.88 | 0.88 | 127 |
| ***Glandular_secretory*** | 0.95 | 0.9 | 0.93 | 124 |
| ***Glandular_secretory_FGF7*** | 1 | 0.98 | 0.99 | 121 |
| ***HOXA13*** | 0.96 | 0.99 | 0.98 | 126 |
| ***Immune_Lymphoid*** | 0.98 | 1 | 0.99 | 131 |
| ***Immune_Myeloid*** | 1 | 0.99 | 0.99 | 138 |
| ***KRT5*** | 0.98 | 0.97 | 0.98 | 126 |
| ***Luminal*** | 0.9 | 0.91 | 0.91 | 123 |

| | | | | |
|---|---|---|---|---|
| ***Lymphatic*** | 1 | 0.99 | 1 | 102 |
| ***MUC5B*** | 0.99 | 0.99 | 0.99 | 119 |
| ***SOX9_basalis*** | 0.97 | 1 | 0.99 | 77 |
| ***SOX9_functionalis_I*** | 0.89 | 0.85 | 0.87 | 128 |
| ***SOX9_functionalis_II*** | 0.82 | 0.87 | 0.84 | 141 |
| ***SOX9_luminal*** | 0.87 | 0.83 | 0.84 | 132 |
| ***Venous*** | 0.91 | 0.93 | 0.92 | 110 |
| ***dHormones*** | 0.9 | 1 | 0.95 | 43 |
| ***dStromal_early*** | 0.79 | 0.82 | 0.81 | 137 |
| ***dStromal_late*** | 0.88 | 0.88 | 0.88 | 120 |
| ***dStromal_mid*** | 0.77 | 0.85 | 0.8 | 124 |
| ***eHormones*** | 0.88 | 0.88 | 0.88 | 129 |
| ***ePV_1a*** | 0.91 | 0.93 | 0.92 | 110 |
| ***ePV_1b*** | 0.93 | 0.93 | 0.93 | 95 |
| ***ePV_2*** | 0.93 | 0.87 | 0.9 | 135 |
| ***eStromal*** | 0.71 | 0.82 | 0.76 | 119 |
| ***eStromal_MMPs*** | 0.99 | 0.89 | 0.94 | 127 |
| ***eStromal_cycling*** | 0.91 | 0.88 | 0.89 | 133 |
| ***mPV*** | 0.96 | 0.94 | 0.95 | 124 |
| ***preCiliated*** | 0.95 | 0.97 | 0.96 | 110 |
| ***preGlandular*** | 0.81 | 0.9 | 0.86 | 135 |
| ***preLuminal*** | 0.87 | 0.92 | 0.9 | 127 |
| ***sHormones*** | 0.92 | 0.88 | 0.9 | 129 |
| ***uSMCs*** | 0.99 | 0.95 | 0.97 | 123 |
| ***Macro average*** | **0.91** | **0.91** | **0.91** | **4328** |
| ***Weighted average*** | **0.91** | **0.91** | **0.91** | **4328** |

**Supplementary Table 3. Mapping of HECA annotations to harmonised major cell types used in this study.** HECA cell types were grouped into broader cell categories to facilitate downstream analyses. For entries labelled “NA”, the original HECA annotations were retained due to insufficient evidence for merging or ambiguity in classification.

| *Main cell type* | *HECA cell type* | *Cell no* |
|---|---|---|
| ***Endothelial*** | Arterial, Lymphatic, Venous | 51,647 |
| ***Epithelial*** | (pre)Ciliated, (pre)glandular, (pre)luminal | 30,707 |
| ***Fibroblasts*** | Fibroblasts basalis | 34,522 |
| ***Lymphoid*** | Immune lymphoid | 119,466 |
| ***Myeloid*** | Immune myeloid | 32,535 |
| ***Perivascular*** | ePV, mPV | 47,531 |
| ***Smooth muscle cells*** | uSMCs | 8,781 |
| **SOX9 basalis** | *SOX9 basalis* | 1,049 |
| ***Stromal*** | dStromal, eStromal | 163,113 |
| ***Stem*** | - | 16,814 |
| ***NA*** | Cycling, Hormones, *HOXA13*, *KRT5*, *MUC5B*, *SOX9* functionalis, *SOX9* luminal | - |

**Supplementary Table 4. Tissues included in each location category for differential expression analysis.**

| *Main location* | *Original sample group* |
|---|---|
| ***Ectopic peritoneum*** | Ectopic endometrium, ectopic adjacent |
| ***Endometrium*** | Eutopic endometrium |
| ***Ovary*** | Ectopic Ovary, unaffected ovary |

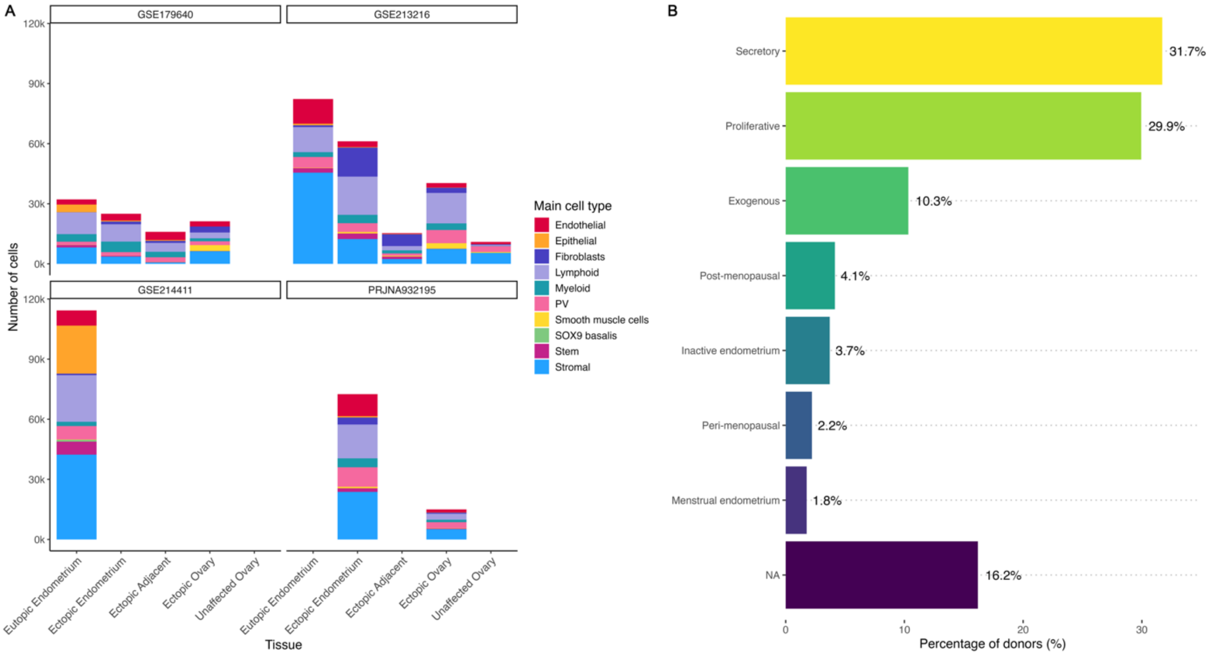


**Supplementary Figure 1. Overview of cell-type composition and donor menstrual cycle stage representation across the datasets. A)** Distribution of main cell types across the datasets and tissues. **B)** Percentage of donors with reported menstrual cycle stages. All NA cases come from the PRJNA932195 dataset. For the analyses we dropped all levels but proliferative, secretory, and exogenous due to small sample sizes.

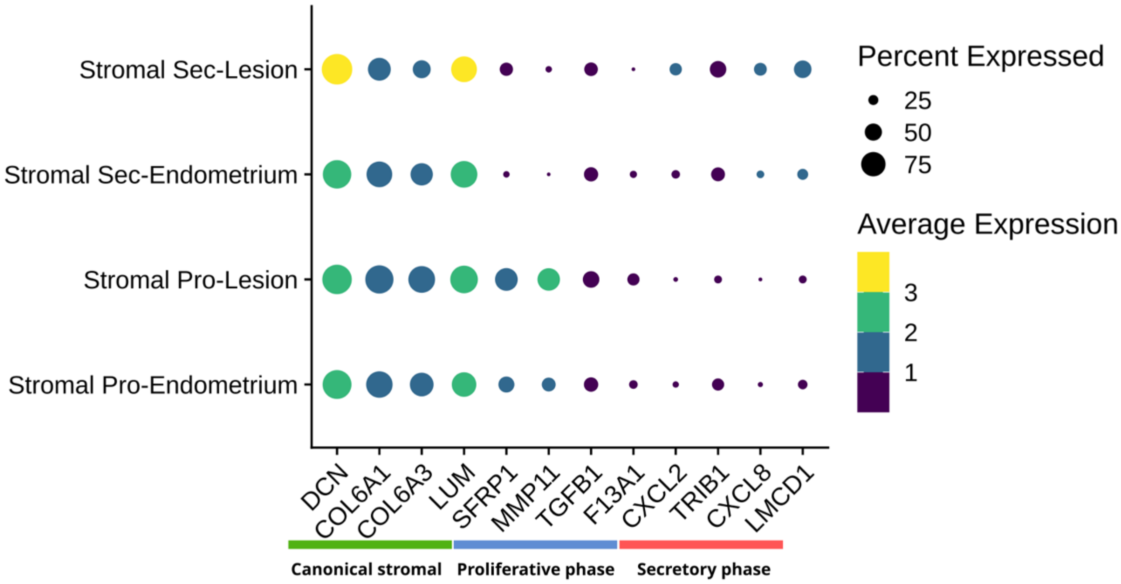


**Supplementary Figure 2. Dot plot displaying the canonical markers.** The canonical markers displayed are for stromal populations, menstrual proliferative and secretory phases in the stromal populations located in endometrium and peritoneal lesions across menstrual cycle phases.
